# FANCJ DNA helicase is recruited to the replisome by AND-1 to ensure genome stability

**DOI:** 10.1101/2022.10.21.513143

**Authors:** Ana Boavida, Luisa M. R. Napolitano, Diana Santos, Giuseppe Cortone, Silvia Onesti, Nanda K. Jegadesan, Dana Branzei, Francesca M. Pisani

## Abstract

FANCJ is a DNA helicase linked to Fanconi anemia and frequently mutated in breast and ovarian cancers. If and how FANCJ is recruited to the replisome is unknown. Here, we report that FANCJ directly binds to AND-1 (the vertebrate ortholog of budding yeast Ctf4), a homo-trimeric protein adaptor that connects the CDC45/MCM2-7/GINS replicative DNA helicase with DNA polymerase α and several factors at DNA replication forks. We find that the interaction between FANCJ and AND-1 requires the integrity of an evolutionarily conserved Ctf4-interacting protein (CIP) box located between the FANCJ helicase motifs IV and V. Disruption of the FANCJ CIP box significantly reduces FANCJ association with the replisome, causing enhanced DNA damage, decreased replication fork recovery and fork asymmetry in stressful conditions. Cancer-relevant FANCJ CIP box variants display reduced AND-1-binding, a finding that suggests a potential role of the mutated *FANCJ* alleles in cancer predisposition.

## Introduction

Faithful DNA replication is critical for genomic stability maintenance. The DNA replication machinery is continuously challenged by damaged DNA templates and other obstacles (alternate secondary structures, R-loops, tightly bound proteins) present throughout the genome. All these physical barriers, which impede a smooth progression of the replication forks, give rise to the so-called replication stress (Saxena & Zou, 2022). This condition is also caused by nucleotide pool depletion or imbalance of replicative factor levels. DNA helicases are important players in counteracting replication stress due to their ability to resolve DNA secondary structures and/or remodel nucleic acid molecules during DNA repair/recombination reactions. The importance of these enzymes in maintaining genome homeostasis is proven by the fact that many of them are genetically linked to rare hereditary diseases characterized by genome instability, chromosome anomalies, developmental defects and cancer predisposition.

FANCJ, also known as BRIP1 (for BRCA1-interacting protein 1), is a super-family 2 (SF2) Iron-Sulphur (Fe-S) cluster-containing DNA helicase, frequently mutated in breast and ovarian cancers, as well as in many other tumor types (Brosh & Cantor, 2014; Cantor *et al*, 2001). FANCJ belongs to the Fanconi anemia pathway and bi-allelic mutations of the encoding gene are known to cause the disease, which is characterized by hematopoietic stem cell defects, progressive bone marrow failure, genomic instability and cancer predisposition (Bridge *et al*, 2005; Levran *et al*, 2005; Levitus *et al*, 2005; Litman *et al*, 2005). Human FANCJ-deficient cell lines display increased sensitivity to Mitomycin C (MMC), a genotoxic agent that introduces DNA inter-strand cross-links (ICLs). FANCJ helicase activity and direct interaction with the mismatch repair protein, MLH1, are both required for processing ICLs (Peng *et al*, 2007). Nonetheless, the precise role of FANCJ in DNA ICL repair reactions remains poorly defined. In addition to BRCA1 and MLH1, other identified FANCJ-binding partners include the DNA exo/endonuclease MRE11 (Suhasini *et al*, 2013), the single-stranded DNA-binding replication protein A (RPA) (Sommers *et al*, 2014), the Bloom DNA helicase (BLM) (Suhasini *et al*, 2011), the trans-lesion synthesis DNA polymerase REV1 (Lowran *et al*, 2019), the Topoisomerase II-binding protein TOPBP1 (Gong *et al*, 2010) and the DNA-end processing nuclease CtIP (Nath & Nagaraju, 2020). These multiple interactions reveal the involvement of FANCJ in various DNA repair pathways as well as in S-phase checkpoint activation. Biochemical studies showed that the purified recombinant FANCJ protein has an ATPase-dependent DNA helicase activity with a 5’ to 3’ directionality that can unwind different DNA substrates *in vitro*, including DNA duplexes forming a fork or containing a 5’-flap, three-stranded displacement loops (D-loops) and various kinds of G-quadruplex (G4) DNA structures (Gupta *et al*, 2005; London *et al*, 2008; Wu & Brosh Jr., 2009). Studies carried out in different systems (worms (Cheung *et al*, 2002), chicken DT40 (Sarkies *et al*, 2012) and human cells (Bharti *et al*, 2013) and *Xenopus laevis* cell-free egg extracts (Castillo Bosch *et al*, 2014; Sato *et al*, 2021)) pointed towards a prominent role of FANCJ in G4 DNA cellular metabolism. Besides, *FANCJ*-knockout (KO) mouse embryonic fibroblasts show increased micro-satellite instability, a phenotype that is further exacerbated upon treatment with compounds that induce replication stress (Matsuzaki *et al*, 2015; Barthelemy *et al*, 2016). All the above considered, it was proposed that FANCJ plays a key role in dismantling DNA secondary structures and unconventional conformations (such as G4) that occur in certain sequence contexts, especially in transiently exposed single-stranded regions. Thus, FANCJ would prevent DNA double-strand break formation and ensure a smooth progression of the DNA replication machineries in stressful conditions (Brosh & Wu, 2021). Notwithstanding the biological significance of FANCJ association with the replication machinery, the protein factor responsible for recruiting FANCJ at the replication forks has not been identified thus far.

Human acidic nucleoplasmic DNA-binding protein 1 (AND-1), also known as WD repeat and high mobility group (HMG)-box DNA-binding protein 1 (WDHD1), is an adaptor protein, crucial for DNA replication, with orthologs in metazoans and in fungi. It was originally identified in a *Saccharomyces cerevisiae* genetic screening of mutants with increased rate of chromosome loss and was named Ctf4 for chromosome transmission fidelity 4 (Spencer *et al*, 1990). Both, budding yeast Ctf4 and human AND-1, were demonstrated to bridge the CDC45/MCM2-7/GINS (CMG) replicative DNA helicase complex, with DNA polymerase α, within the replication machinery (Gambus *et al*, 2009; Tanaka *et al*, 2009; Simon *et al*, 2014; Kilkenny *et al*, 2017; Guan *et al*, 2017). In budding yeast, Ctf4 loss has pleiotropic effects, including increased sensitivity to genotoxic agents, defective sister chromatid cohesion and modifications in the ribosomal DNA genomic *locus* (Villa *et al*, 2016; Samora *et al*, 2016; Fumasoni *et al*, 2015; Sasaki & Kobayashi, 2017). Instead, the Ctf4 orthologous proteins from *Schizosaccharomyces pombe* (Williams & McIntosh, 2005), *Drosophila* (Gosnell & Christensen, 2011) and chicken (DT40 cell system) (Abe *et al*, 2018) are essential for cell proliferation. In human cells, AND-1 depletion using siRNAs slows down cell cycle progression (Yoshizawa-Sugata & Masai, 2009). Subsequent structural studies revealed that budding yeast Ctf4 and human AND-1 share a similar multi-domain organization, each comprising a β-propeller (WD40) and a SepB module (Simon *et al*, 2014; Kilkenny *et al*, 2017; Guan *et al*, 2017). However, the human AND-1 polypeptide chain has a C-terminal extension that includes a HMG DNA-binding domain, not present in budding yeast Ctf4. The SepB domain is responsible for AND-1/Ctf4 trimerization and for interaction with proteins containing the so-called Ctf4-interacting protein (CIP) box. This is a conserved short peptide sequence, originally identified in *S. cerevisiae* DNA polymerase α catalytic subunit and in GINS Sld5, which contains conserved acidic and hydrophobic amino-acid residues essential for binding Ctf4 (Simon *et al*, 2014). Other Ctf4-binding partners were identified in budding yeast, such as the helicase–nuclease Dna2, the ribosomal DNA compaction protein Tof2, the replication initiation factor Dpb2 and the sister chromatid cohesion helicase Chl1 (Simon *et al*, 2014; Villa *et al*, 2016; Samora *et al*, 2016). These findings suggest that Ctf4 is an interaction hub connecting the DNA replication machinery to multiple proteins and enzymes that contribute to diverse aspects of genome duplication.

Here, we report that human FANCJ directly interacts with AND-1 through a conserved CIP box that lies between the DNA helicase motifs IV and V. FANCJ/AND-1 interaction is critical for recruiting FANCJ to the replisome and for promoting a smooth progression of the DNA replication forks. We show that faulty association of FANCJ to AND-1, due to CIP box mutations, gives rise to increased DNA damage, replication fork stalling and asymmetry in either unperturbed or stressful conditions. Besides, we report evidence that cancer-associated *FANCJ* mutant alleles causing substitutions of CIP box amino-acid residues are defective in AND-1-binding. These findings about the functional consequences of clinically relevant *FANCJ* missense variants provide important clues on how to interpret cancer risk and devise novel therapeutic strategies.

## Results

### Identification of a AND-1/Ctf4-interacting protein (CIP) box in FANCJ

The *S. cerevisiae* Chl1 DNA helicase was demonstrated to be recruited to the DNA replication forks through a direct interaction with Ctf4. This interaction requires the integrity of a CIP box that is located between the conserved helicase motifs IV and V of the Chl1 polypeptide chain (Samora *et al*, 2016). Human DDX11 and FANCJ are paralogous proteins, both sharing sequence similarity with budding yeast Chl1. A bioinformatic analysis revealed the presence of a putative CIP box in the human FANCJ polypeptide chain between the helicase motifs IV and V (see a multiple sequence alignment of the CIP box from various Ctf4/AND-1 client proteins in Figure 1A). Besides, a putative CIP box is invariantly present in the same position of many vertebrate FANCJ orthologs, and amino-acid residues reported to be critical for AND-1/Ctf4- binding (Villa *et al*, 2016; Guan *et al*, 2017; Kilkenny *et al*, 2017) appeared to be highly conserved in the aligned FANCJ sequences (Figure 7A). A model of the human FANCJ three-dimensional structure, based on artificial intelligence, reveals that the putative CIP box folds as an α-helix (Figure 1B). These findings prompted us to test if human FANCJ interacts with AND-1 and if this interaction is mediated by the newly identified putative CIP box. Hence, we substituted highly conserved amino-acid residues of the CIP box with Alanine to generate a FANCJ mutant that was named FANCJ AALA (Figure 1A). HEK 293T cells were transiently transfected with plasmid vectors expressing Flag-tagged FANCJ wild type (WT) or the AALA mutant (AALA) and co-immunoprecipitation experiments with anti-Flag M2 beads in whole cell extracts were performed. The results revealed that ectopically expressed Flag-tagged FANCJ WT was co-pulled-down with the endogenous AND-1 protein (Figure 1C). In contrast, the amount of AND-1 co-immuno-precipitated with the Flag-tagged FANCJ AALA mutant was reduced by about 3-fold (Figure 1C). These results suggest that FANCJ associates with AND-1 in cell extracts and their association is mainly dependent on the integrity of the newly identified FANCJ CIP box.

**Figure 1.**
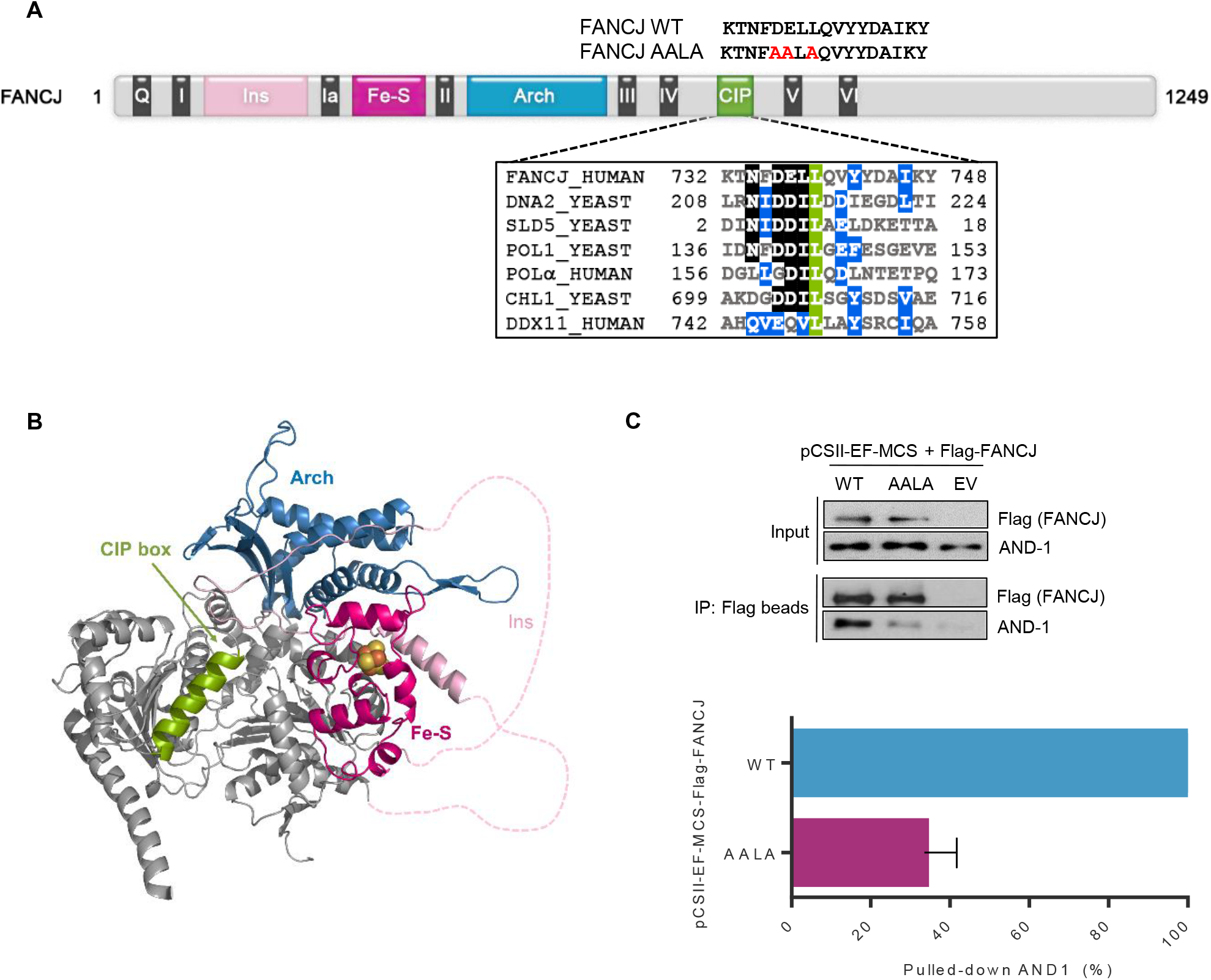
Identification of an AND-1/Ctf4-interacting protein (CIP) box in FANCJ. **A.** Schematic representation of the polypeptide chain of human FANCJ. Conserved helicase motifs (from I to VI) are indicated in *dark grey* (for simplicity motifs Va and Vb are not shown). The identified CIP box is depicted in *green*. Other sequence motifs and specific domains are indicated with different colors, using the abbreviations: *Q*, for Q motif; *Ins*, for Insertion; *Fe-S*, for Fe-S cluster; *Arch*, for Arch domain. A multiple sequence alignment of human FANCJ and other Ctf4-/AND-1-interacting proteins CIP boxes is reported in the inset. The KALIGN tool (version 3.3.1) was used. Invariant, identical and similar residues are highlighted in *green*, *black* and *blue*, respectively. The following abbreviations are used: *HUMAN*, *Homo sapiens*; *YEAST*, *Saccharomyces cerevisiae*; *CHL1*, chromosome loss 1; *POL1* and *Polα*, DNA polymerase α catalytic subunit; *SLD5*, synthetic lethality with *dpb11-1* protein 5. CIP box mutations to generate the FANCJ AALA mutant are indicated in *red* above the diagram. **B.** Homology model of the FANCJ protein, as predicted by the AlphaFold server (Jumper *et al*, 2021) illustrating the location of the indicated conserved domains. Color code is the same as in panel *a*. Disordered regions within the insertion are indicated by dashed lines; the Fe-S cluster was manually modelled based on the crystal structures of homologous helicases. **C.** Co-immunoprecipitation experiments were carried out with anti-Flag M2 agarose beads on whole extracts from HEK 293T cells transfected with pCSII-EF-MCS vector empty (*EV*) or expressing Flag-tagged FANCJ wild type (*WT*) or the AALA mutant (*AALA*). Western blot analysis is shown, where the endogenous AND-1 protein was detected using specific antibodies. Co-immunoprecipitation experiments were done in triplicate. Level of immuno-precipitated AND-1 was normalized to pulled down Flag-tagged FANCJ in each sample. *Error bars* represent standard deviation.

Then, we purified Flag-tagged FANCJ WT and AALA mutant proteins from transiently transfected HEK 293T cells and measured their DNA helicase activity using a fluorescent-labelled DNA substrate in a gel-based assay. We found that the FANCJ AALA mutant retained a level of DNA unwinding activity comparable with that one of FANCJ WT either on a G4 DNA substrate or on a forked duplex (Figure 7D and Appendix Figure S3), ruling out the possibility that the CIP box mutations could have destabilized the recombinant protein and thus the loss of binding affinity was simply due to misfolding. Moreover, these results indicate that the FANCJ AALA variant is a separation-of-function mutant, retaining DNA helicase activity, but being almost completely unable to associate with AND-1 in cell extracts.

### FANCJ directly interacts with the AND-1 SepB domain via its CIP box

To examine whether the interaction between FANCJ and AND-1 was direct, *in vitro* co-pull-down assays were carried out. In these experiments we used the His-tagged SepB domain of human AND-1 produced in bacterial cells as a soluble homo-trimeric complex (6xHis SepB) (Kilkenny *et al*, 2017; Guan *et al*, 2017). Biochemical and structural studies revealed that human AND-1 and yeast Ctf4 share a similar three-dimensional structure, formed by a N-terminal WD40 β-propeller domain and a SepB module (Figure 2A). The latter is responsible for AND-1/Ctf4 trimerization and for binding CIP box-containing proteins. We found that FANCJ WT was able to pull-down 6xHis-SepB, while binding by FANCJ AALA mutant was almost completely abrogated (Figure 2B). We next expressed a short peptide containing the FANCJ CIP-box or its AALA mutant version as fusion with the GST protein. We found that the AND-1 SepB domain was pulled-down by the GST-FANCJ CIP box WT chimeric protein but not by the AALA mutant (Figure 2C). Structural studies revealed that the SepB domain of either human AND-1 or budding yeast Ctf4 is made of a β-propeller subdomain followed by a bundle of 5 α-helixes (Simon *et al*, 2014; Kilkenny *et al*, 2017; Guan *et al*, 2017). The binding surface for the AND-1/Ctf4 client proteins consists of a groove between α-helixes 2 and 4 (H2 and H4; Figure 2D) of the SepB helical bundle. It was reported that the side chain of Methionine766, which belongs to α-helix H2, is exposed on the groove surface and its substitution with an Alanine residue almost totally abolishes the interaction of the SepB domain with the catalytic subunit of human DNA polymerase α, an AND-1 client protein (Guan *et al*, 2017; Kilkenny *et al*, 2017). To identify the AND-1 structural elements responsible for FANCJ CIP box-binding, we produced the SepB α-helix bundle and its Met766Ala mutant derivative as GST-fusion proteins (named GST-SepB α-helix bundle WT and M766A, respectively) and used these purified chimeric proteins to pull-down FANCJ WT and AALA mutant. As shown in Figure 2E, a direct interaction was observed only between GST-SepB α-helix bundle WT (not the M766A mutant) and FANCJ WT (not the AALA mutant). Collectively, these data demonstrate that FANCJ directly binds to AND-1 and this interaction requires the integrity of either the FANCJ CIP box and the AND-1 SepB α-helix bundle.

**Figure 2.**
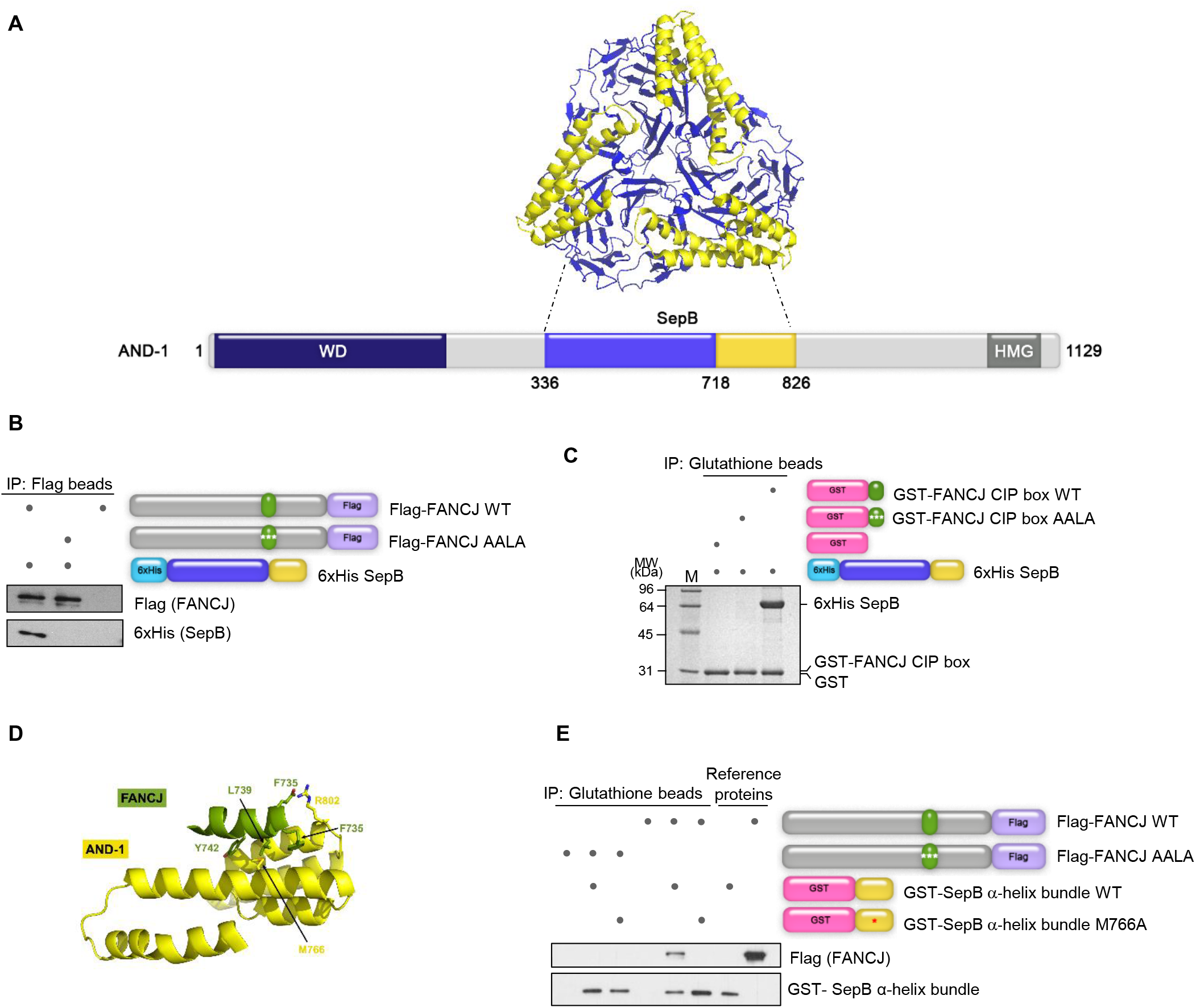
FANCJ CIP box directly binds the AND-1 SepB domain. **A.** Schematic representation of the polypeptide chain of human AND-1. The WD 40 repeat (*WD*), SepB (*SepB*) and high mobility group (*HMG*) domains are indicated by boxes of different colors. The β-propeller subdomain and α-helix bundle that form the SepB domain are depicted in *blue* and *yellow*, respectively. Same colors are used to highlight them in the human AND-1 SepB structure (Protein Data Bank ID 5gvb). **B**. Co-pull-down experiments of Flag-tagged FANCJ WT or AALA mutant and AND-1 6xHis-tagged SepB using anti-Flag M2 agarose beads. Pulled-down samples were analyzed by Western blot with the indicated antibodies. **C**. GST was fused to the FANCJ CIP box (amino acid residues 730-747) for use in a pull-down assay with purified AND-1 6xHis SepB. Specific mutations of the FANCJ CIP box (as in the GST-FANCJ CIP box AALA derivative) disrupt its ability to pull-down the SepB domain. **D**. A structural model of the interaction surface between the FANCJ CIP box and the AND-1 SepB α-helix bundle; a model was built, manually adjusted and subjected to energy optimization, by assembling the AND-1 crystal structure (PDB ID: 5gvb) with a model prediction for the FANCJ CIP box helix, based on the crystal structure of budding yeast Ctf4/Pol1-CIP (PDB ID: 4c93). Amino-acid residues side chains, putatively involved in critical contacts, are shown as *sticks*. **E**. The GST-SepB α-helix bundle chimeric protein was able to pull down purified Flag-tagged FANCJ WT. Substitution of the critical Met766 with Ala in the SepB α-helix bundle abolishes FANCJ-binding.

### Establishment of FANCJ-knock-out and complementation by FANCJ wild type and AALA mutant alleles

To examine the physiological relevance of the FANCJ/AND-1 interaction, we carried out complementation assays in cells lacking the endogenous FANCJ protein and ectopically expressing *FANCJ* WT allele or its AALA mutant derivative. To this end, we generated a HeLa cell line where *FANCJ* was knocked-out (FJ-KO) using CRISPR-paired guide RNAs targeting exons 2 and 3 (Figure 3A). Western blot analysis and Sanger sequencing of selected clones were carried out to confirm FANCJ protein loss and identify specific mutations introduced by the Cas9 activity (Figure 3C). Since FANCJ depletion is expected to sensitize cells to Mitomycin C (MMC), a DNA inter-strand cross-linking agent, and Pyridostatin (PDS), a G4 DNA stabilizer, we examined the viability of the FJ-KO HeLa cells after 5-day treatment with increasing doses of these drugs. As shown in Figure 3B, the FJ-KO HeLa cell line is sensitive to either MMC or PDS, with MMC exerting a toxic effect even in a low nM concentration range (10-20 nM). Afterwards, we stably complemented the FJ-KO HeLa line by expressing Flag-tagged FANCJ WT or the AALA mutant under the control of a Tetracycline-responsive promoter using the pMK240 plasmid vector (Okumura *et al*, 2018). Western blot analyses revealed that FANCJ WT and the AALA mutant were expressed at a comparable level and only upon Doxycycline induction in the selected clones (Figure 3C). These stably complemented cell lines were used to analyze the impact of FANCJ/AND-1 interaction on genome stability maintenance.

**Figure 3.**
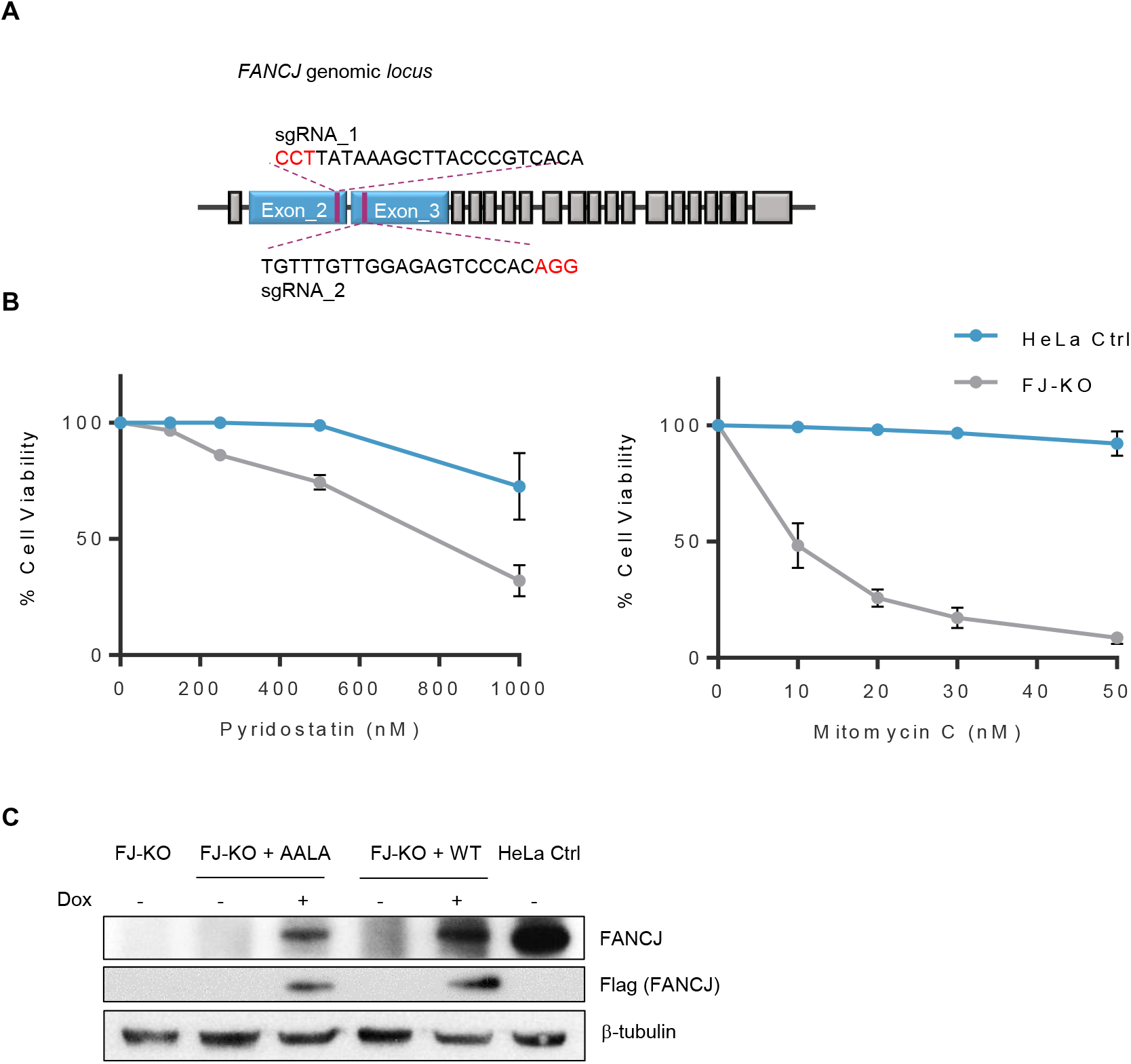
Establishment of a FJ-KO HeLa cell line and its complementation by *FANCJ* WT and AALA mutant alleles. **A.** Schematic representation of the human *FANCJ* genomic *locus*. The sequence of CRISPR-paired guide RNAs targeting exons 2 and 3 is reported. PAM sequence is highlighted in *red*. **B.** Cell viability assays of wild type control (*Ctrl*) and FJ-KO HeLa cell lines treated for 5 days with the indicated concentrations of MMC and PDS (n = 3 biologically independent experiments, mean ± SD). **C**. FJ-KO HeLa cell lines were established that stably express Flag-tagged FANCJ WT or the AALA mutant under the control of a Tetracycline-responsive promoter. Expression of ectopic *FANCJ* was detected before and after induction with Doxycycline (1 μg/mL; *Dox*) by Western blot analysis of whole extracts from the indicated cell lines using an anti-FANCJ or anti-Flag antibody. β-tubulin was used as a loading control.

### FANCJ loss or its reduced AND-1-binding enhances DNA damage

Thereafter, we examined whether the observed reduced viability of the FJ-KO HeLa cell line upon treatment with MMC or PDS was associated with increased DNA damage. After verifying the expression of the Flag-FANCJ WT and AALA mutant upon Doxycycline induction with or without drug treatment (Appendix Figure S2), we analyzed formation of γ-H2A.X *foci* by indirect immuno-fluorescence experiments. The level of DNA damage was higher in FJ-KO compared to HeLa control cells, not only following treatment with MMC and PDS, but also in unperturbed conditions (Figure 4). Besides, complementation with the *FANCJ* WT allele almost completely reverted the above phenotype, while expression of the *FANCJ* AALA variant gave rise to a level of γ-H2A.X *foci* even higher compared to the FJ-KO cells (either with or without drug treatment; Figure 4) suggesting a possible dominant negative behavior of the *FANCJ* AALA allele. These results indicated that the FANCJ/AND-1 interaction is critical for suppressing DNA damage either in unchallenged conditions or after inducing DNA ICLs (treatment with MMC) or stabilizing G4 DNA structures (treatment with PDS).

**Figure 4.**
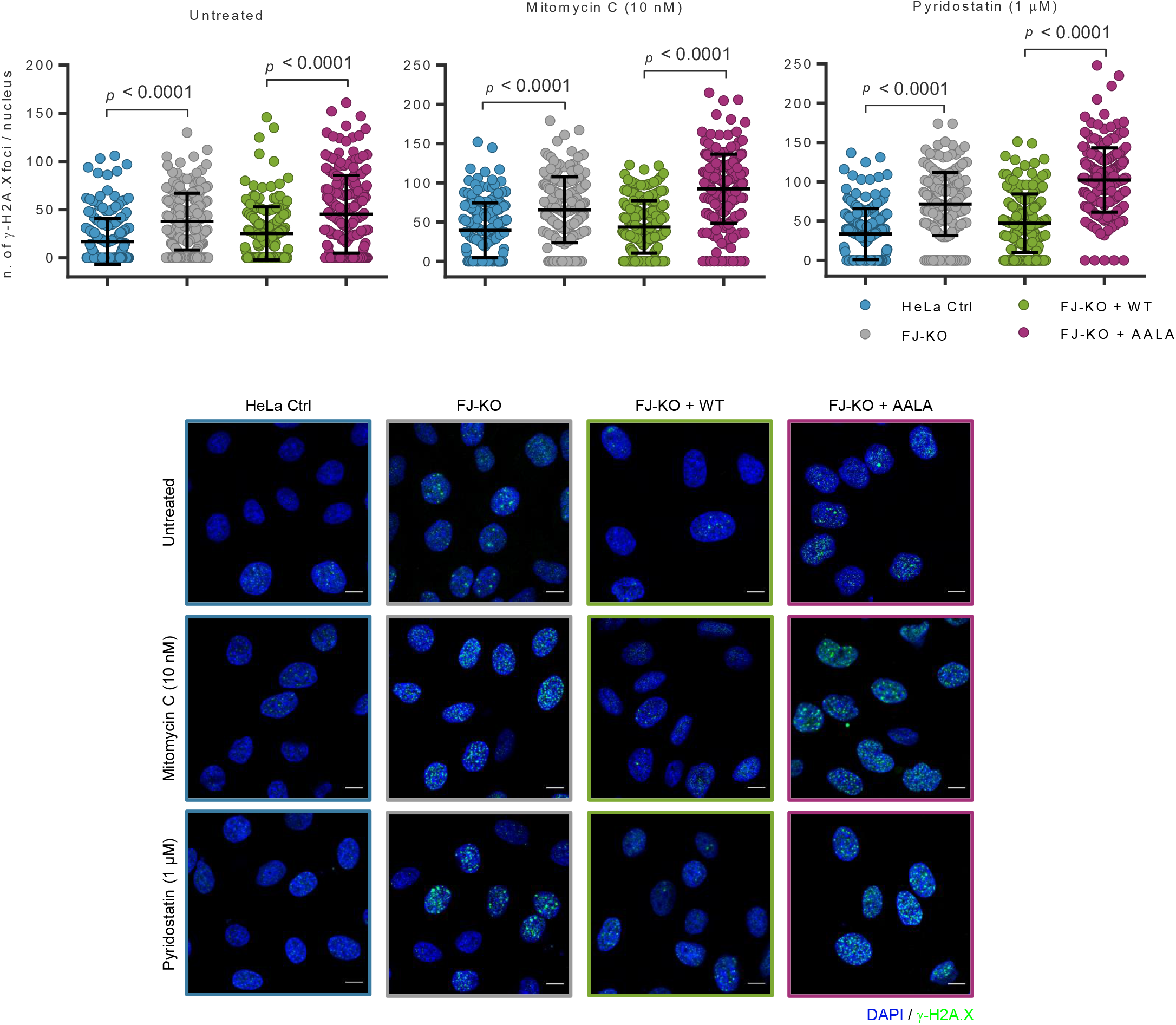
FANCJ loss or reduced binding to AND-1 enhance DNA damage. The indicated cell lines were treated with PDS or MMC, after Doxycycline induction. γ-H2A.X *focus* formation was detected by immunofluorescence with a monoclonal antibody that specifically recognizes the Ser139-phosphorylated form of the above histone. Scale bar, 10 μM. Dot plot of the number of *foci* detected per cell is reported in each graph. *Bars* indicate mean ± SD; 200 cells were analyzed per condition; *n* = 2 biologically independent experiments, with at least 2 technical replicates per experiment; *p*-value (*p* < 0.01) was calculated using Student’s *t*-test non-parametric for unpaired data with Welch correction.

### FANCJ associates with the DNA replication machinery through a direct interaction with AND-1

Given that FANCJ directly binds to AND-1 *via* its CIP box and AND-1 is a component of the DNA replication machinery, we then asked whether FANCJ is localized at active DNA replication forks. Co-immunoprecipitation experiments were carried out to identify proteins associated with CDC45, a component of the CMG complex in cell extracts. HEK 293T cells, transiently transfected with plasmids expressing Flag-tagged FANCJ WT or the AALA mutant, were released in S phase after an overnight thymidine block. The nuclear fraction was prepared from synchronized cells using a procedure that included a sonication step followed by an exhaustive nuclease treatment to remove nucleic acids that could mediate protein-protein interactions. Co-immunoprecipitations were performed in the nuclear fraction samples with an anti-CDC45 antibody conjugated to Protein A Sepharose beads. As shown in Figure 5A, Western blot analyses revealed that FANCJ WT was pulled-down with CDC45, together with AND-1 and MCM4, a subunit of the MCM2-7 complex; in contrast, the amount of FANCJ AALA mutant co-immunoprecipitated with CDC45 was reduced by approximately 3-fold in experiment replicates. Since CDC45 is loaded onto chromatin only in S phase as a stable component of the CMG complex (Masai *et al*, 2010), our results revealed that FANCJ was anchored to the replication machinery *via* a direct interaction with AND-1, as this interaction was dependent on the integrity of the FANCJ CIP box. Thereafter, we examined localization of FANCJ to DNA synthesis sites marked by EdU incorporation in FJ-KO HeLa cell lines complemented with *FANCJ* WT or AALA mutant allele using the *in situ* visualization of protein interactions at DNA replication forks (SiRF) technique (Roy *et al*, 2018). Control experiments revealed a comparable level of EdU-EdU proximity ligation assay (PLA) spots in all the cell lines tested, meaning that they had a similar number of active replication factories and equal probability to produce a positive signal in the conditions used for the SiRF assay (Figure 5B). Our analyses revealed the presence of robust FANCJ-EdU PLA signals either in HeLa control cells or in the FJ-KO line complemented with the *FANCJ* WT allele, indicating that FANCJ localizes to DNA synthesis sites. In contrast, EdU-FANCJ PLA spots were almost totally absent in FJ-KO HeLa cells and FJ-KO line complemented with the FANCJ AALA mutant, suggesting that CIP box mutations remarkably reduced the level of FANCJ associated to DNA replication factories (Figure 5B).

**Figure 5.**
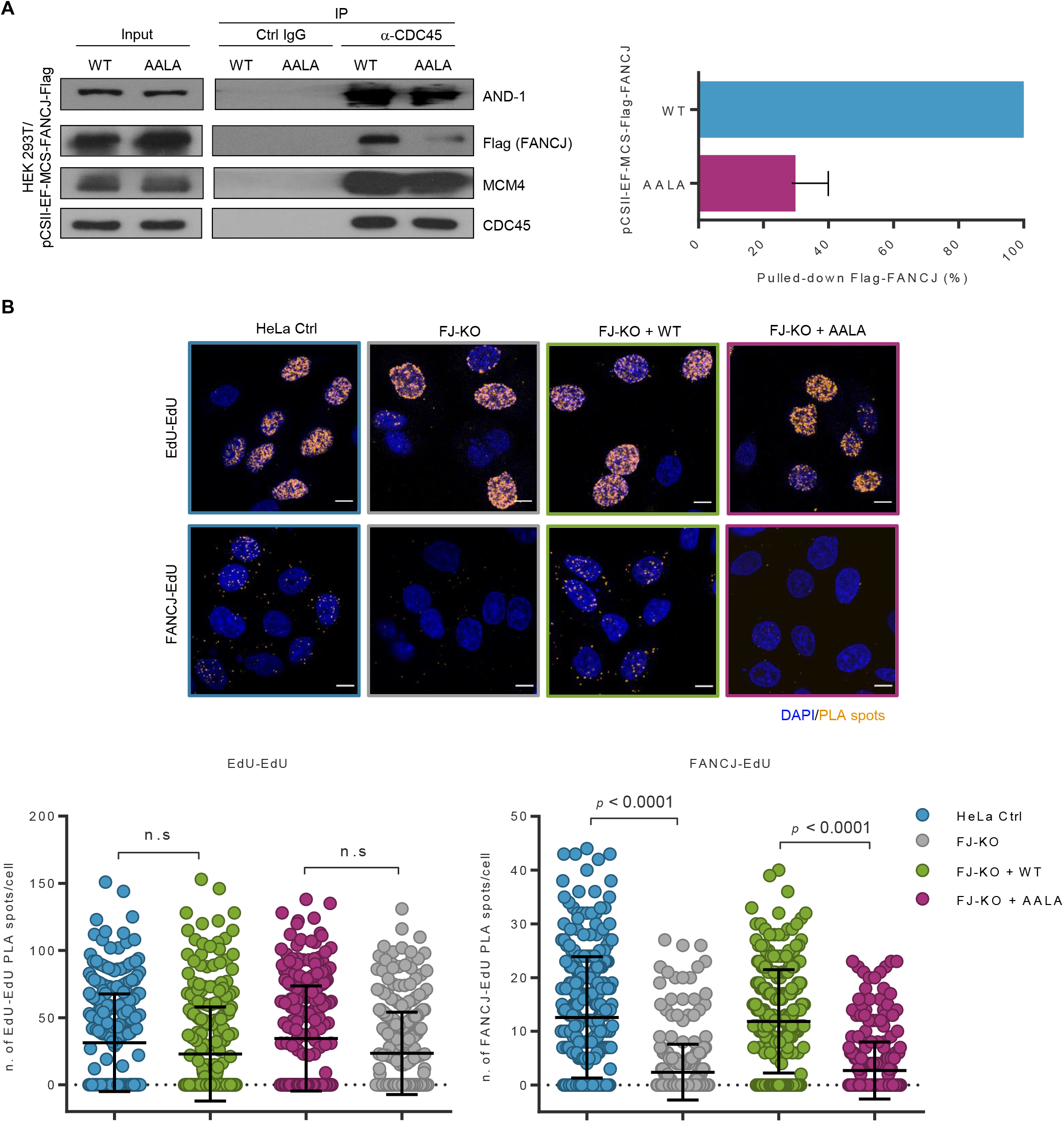
FANCJ associates to DNA replication forks through a direct interaction with AND-1. **A**. Co-immunoprecipitation experiments were carried out in the nuclear fraction of HEK 293T cells transiently transfected with pCSII-EF-MCS plasmid constructs expressing Flag-tagged FANCJ wild type (*WT*) or AALA mutant (*AALA*). Control (*Ctrl-IgG*) and anti-CDC45 rabbit IgG (*α*-CDC45), bound to Protein A Sepharose beads, were added to each indicated nuclear fraction. Immuno-precipitated protein samples were analyzed by Western blot using the indicated antibodies. Data from triplicate experiments show level of immuno-precipitated FANCJ (mean ± SD), normalized to pulled-down endogenous CDC45, in each sample. Means with standard errors are shown. **B**. FANCJ association to sites of DNA synthesis was analyzed by SiRF assays. EdU-EdU (*upper part*) or FANCJ-EdU (*lower part*) proximity ligation assay (PLA) spots were quantified from control (*Ctrl*), FJ-KO and complemented HeLa cell lines. Frequency distribution of population was analyzed and *p*-value (*p* < 0.01) was calculated using Student’s *t*-test non-parametric for unpaired data with Welch correction (< 300 cells per condition; *n* = 3 biologically independent experiments; *bars* show mean ± SD).

Collectively, these results reveal that FANCJ is anchored to the replisomes *via* a direct interaction with AND-1 and FANCJ CIP box mutations almost completely disrupt its co-localization with the DNA replication factories.

### AND-1-binding by FANCJ promotes replication fork progression in challenged conditions

Next, we examined the effect of disrupting the FANCJ CIP box on the replication fork dynamics using DNA fiber track assays to monitor DNA replication at a single molecule level in the aforementioned cell lines. As schematically depicted in Figure 6A, cells were first labelled with CldU and then a second label, IdU, was administered without or with PDS for testing replication fork progression in normal and challenging conditions, respectively. A significant reduction of replication fork speed was measured upon PDS treatment in FJ-KO cells and in FJ-KO cells complemented with the AALA mutant, with a more prominent effect observed in this latter condition (Figure 6A). Analysis of sister forks (forks emanating from the same replication origin), obtained in PDS-treated conditions, revealed that the percentage of those with asymmetric tracks was remarkably higher in either FJ-KO cells or in FJ-KO cells complemented with the AALA mutant, indicating that replication forks stalled more frequently when FANCJ is either absent or not stably anchored to the ongoing replisomes (Figure 6B).

**Figure 6.**
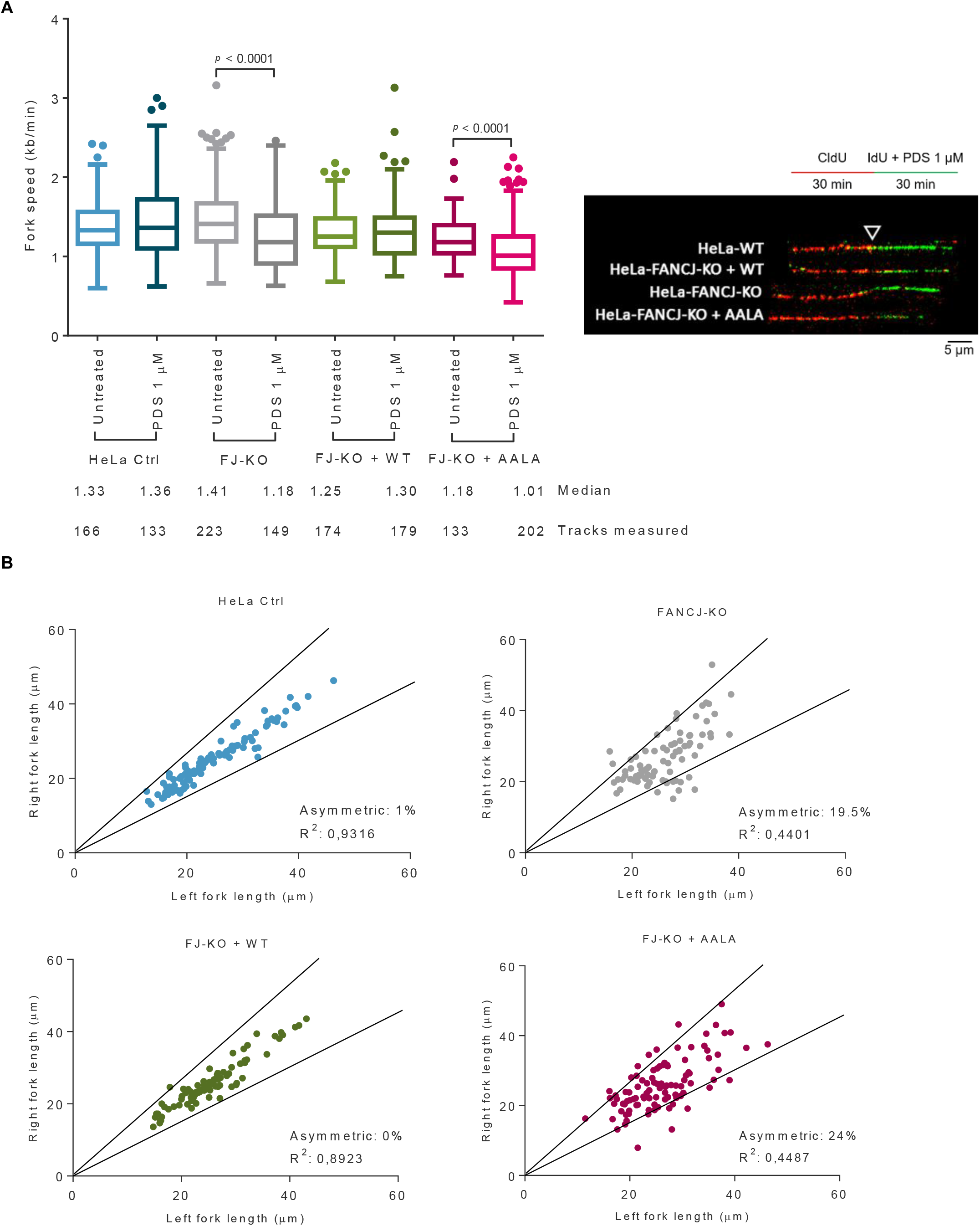
Association of FANCJ to the DNA replication machinery promotes fork progression in challenged conditions. **A**. DNA fiber track analysis was carried out in either untreated or perturbed conditions. Box-plot showing fork speed determined on CldU:IdU double-labeled DNA fibers from control (*Ctrl*), FJ-KO and stably complemented HeLa cell lines, untreated or treated with PDS (1μM) simultaneously with IdU pulse (*n* = 3 biologically independent experiments, with at least 2 technical replicates). Labeled DNA fibers are shown from a representative experiment. Median fork rate and the number of tracks analyzed are shown. The box extends from the 25th minus 1.5IQR to 75th plus 1.5IQR percentiles – Tukey method. *p* values were calculated by Mann–Whitney U test. **B**. Fork asymmetry analysis in cells treated with PDS during IdU pulse, as in panel a. The central area between the lines delimits a variation of < 25% in fiber length. R^2^ is the linear correlation coefficient. Approximately 100 sister forks (green-red-green tracks only) were analyzed and plotted from 3 biologically independent experiments.

**Figure 7.**
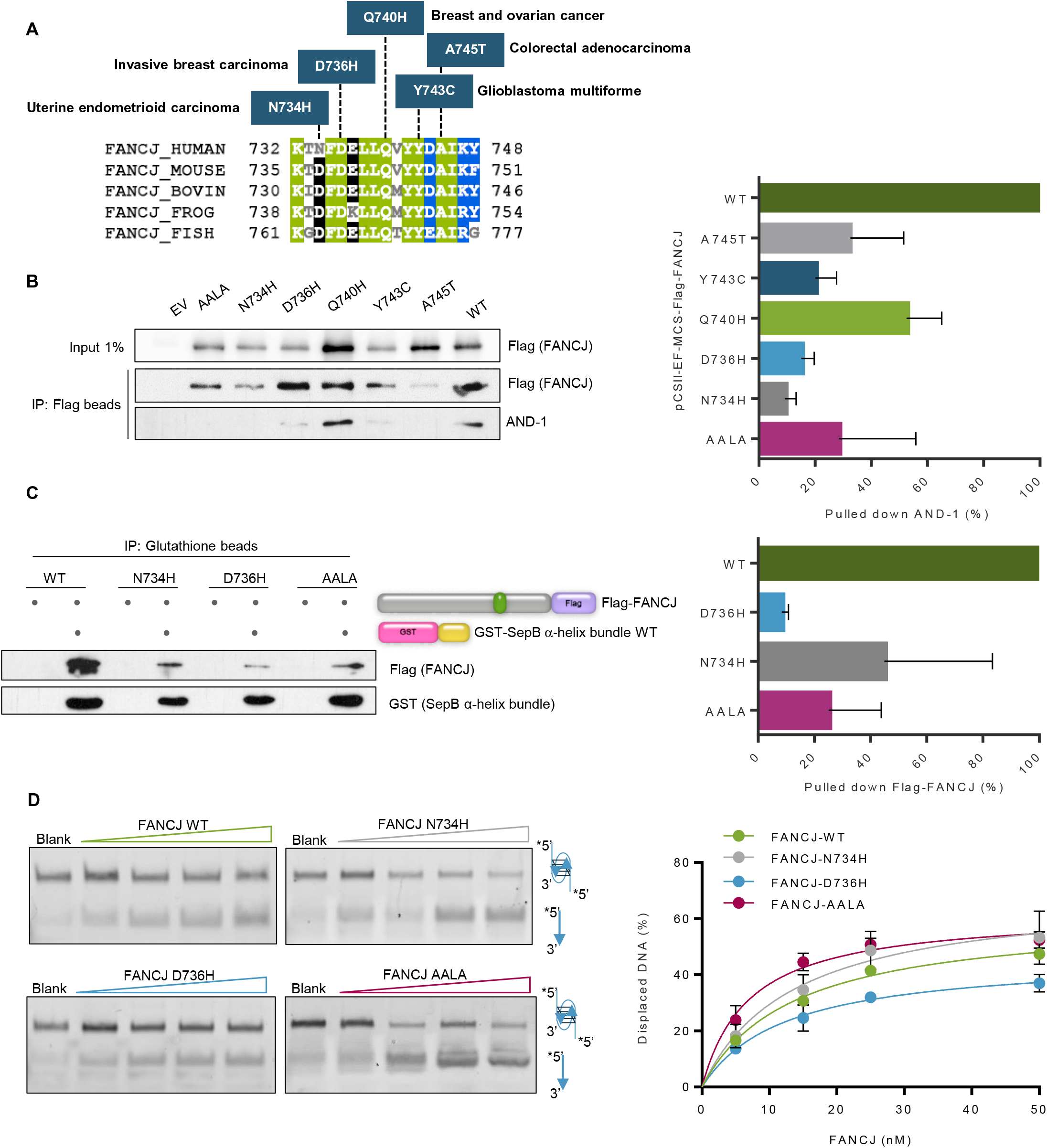
Cancer-associated FANCJ CIP box mutations show reduced AND-1-binding. **A.** Cancer-relevant FANCJ CIP box mutants. Amino-acid changes corresponding to *FANCJ* missense variants found in the indicated tumors are shown in the upper part. A multiple sequence alignment of various FANCJ orthologs CIP box is reported in the lower part; invariant, identical and similar residues are highlighted in *green*, *black* and blue, respectively. The following abbreviations were used: *HUMAN*, Homo sapiens; *MOUSE*, *Mus musculus*; *BOVIN*, *Bos taurus*; *CHICK*, *Gallus gallus*; *FROG*, *Xenopus laevis*; *FISH*, *Danio rerio*. **B.** Co-immunoprecipitation experiments with anti-Flag M2 agarose beads were carried out on nuclear extracts of HeLa FJ-KO cells transiently transfected with pCSII-EF-MCS vector empty (*EV*) or expressing Flag-tagged FANCJ wild type (*WT*) or the indicated mutants (*AALA*, *N734H*, *D736H*, *Q740H*, *Y743C* and *A745T*). Western blot analysis is shown, where the endogenous AND-1 protein was detected using specific antibodies. Co-immunoprecipitation experiments were done in triplicate. Level of immuno-precipitated AND-1 was normalized to pulled-down Flag-tagged FANCJ in each sample. Mean ± SD are shown in the bar graph. **C.** GST-pull-down experiments of GST-SepB α-helix bundle protein and Flag-tagged FANCJ WT, AALA, N734H or D736H mutants. Reduced binding to SepB α-helix bundle is observed for all the FANCJ variants, compared to the wild type protein. Mean ± SD are shown in the bar graph (*n* = 3 biologically independent experiments). **D.** Gel-based DNA helicase assays were carried out using FANCJ WT or the indicated mutants at increasing concentrations and a fluorescent-labeled anti-parallel bi-molecular G4 DNA substrate (*n* = 3 independent experiments, mean ± SD). *Asterisk* represents the fluorophore attached to the DNA oligonucleotide.

### Cancer-relevant FANCJ CIP box mutations reduce AND-1-binding

After *BRCA1* and *BRCA2*, *FANCJ* is the third most common ovarian cancer susceptibility gene: nearly 0.9% - 2.5% of all ovarian cancer patients carry a splice-site, stop or frameshift mutation in the *FANCJ* gene (Ramus *et al*, 2015; Norquist *et al*, 2016). Besides, *FANCJ* was found to be mutated in several other malignancies, including melanoma, breast, prostate, and hereditary colon cancer, suggesting that *FANCJ* mutations may be a risk factor in multiple tumor types (Cantor & Guillemette, 2011; Paulo *et al*, 2018; Ali *et al*, 2019). Nevertheless, a direct association of *FANCJ* mutations with predisposition to breast cancer has been questioned (Weber-Lassalle *et al*, 2018; Easton *et al*, 2016). However, as the majority of *FANCJ* clinical variants remain uncharacterized, their possible connection to cancer risk is difficult to interpret (Moyer *et al*, 2020; Calvo *et al*, 2021). Through data mining of cancer genomics databases (cBioPortal, http://cbioportal.org, Cosmic-3D, https://cancer.sanger.ac.uk/cosmic3d, and gnomAD, https://gnomad.broadinstitute.org) we pinpointed *FANCJ* variants expected to give rise to mutations of CIP box amino-acid residues in various malignancies, including uterine endometrial carcinoma (N734H), invasive breast carcinoma (D736H), breast and ovarian cancer (Q740H), glioblastoma multiforme (Y743C) and colorectal adenocarcinoma (A745T) (Figure 7A). To test AND-1-binding by the above cancer-associated FANCJ mutant derivatives, we carried out co-immunoprecipitations experiments with anti-Flag agarose beads in whole extracts of FJ-KO HeLa cells transiently transfected with plasmids expressing Flag-tagged versions of the FANCJ mutants of interest. As reported in Figure 7B, all the examined cancer-associated FANCJ CIP box mutants displayed a reduced association with the endogenous AND-1 protein. The most dramatic effect was observed with the FANCJ variants N734H and D736H, as their binding to AND-1 in cell extracts was almost completely abrogated. The FANCJ mutants N734H and D736H were produced as Flag-tagged recombinant proteins and purified from transiently transfected HEK 293T cells (Appendix Figure S3A). Their direct interaction with the GST-SepB α-helix bundle fusion protein was examined by GST-pull down assays. Results from these experiments confirmed that AND-1-binding by both these FANCJ mutants was remarkably reduced (Figure 7C). DNA helicase activity assays revealed that the FANCJ N734H and D736H mutants were able to resolve either an anti-parallel bi-molecular G4 or a forked duplex DNA substrate with an efficiency comparable with those of FANCJ wild type and the AALA mutant (Figure 7D and Appendix Figure S3B), confirming separation of function between recruitment to AND-1 and the DNA helicase activity.

## Discussion

Here, we report the identification of an AND-1/Ctf4-interacting protein (CIP) box in the human FANCJ sequence, located between the conserved DNA helicase motifs IV and V. We demonstrated that FANCJ directly binds to the α-helix bundle of the AND-1 SepB domain through this newly identified CIP box. Although a partially conserved CIP box was found to be present also in the corresponding DDX11 sequence, we were unable to detect any direct interaction between Flag-tagged DDX11 and the AND-1 SepB domain in co-immunoprecipitations carried out *in vitro* with the purified recombinant proteins (Appendix Figure S1A-B), in agreement with a previous report (Farina *et al*, 2008). The physiological relevance of the FANCJ/AND-1 interaction was examined by functional assays carried out in FJ-KO HeLa cells stably complemented with either *FANCJ* WT or the AALA mutant allele. The FANCJ AALA derivative is a separation-of-function mutant as it is defective in AND-1-binding, due to changes of critical CIP box residues, but retains a level of DNA helicase activity comparable with the wild type protein on either forked duplex or G4 DNA substrates. Our analyses revealed that the integrity of the CIP box was critical for recruiting FANCJ to the replisome through a direct binding to AND-1 and for suppressing DNA damage in either unperturbed or challenged conditions (treatment with MMC, an ICL agent, or PDS, a G4 DNA-stabilizer). Besides, the FANCJ/AND-1 interaction protected DNA replication forks by promoting their progression and stability, especially in cells treated with PDS. These findings are consistent with several lines of evidence suggesting that FANCJ has a prominent role in resolving G4 structures at DNA replication forks (Cheung *et al*, 2002; Wu *et al*, 2008; Schwab *et al*, 2013; Castillo Bosch *et al*, 2014; Van Schendel *et al*, 2021) and with biochemical studies showing that FANCJ is able to untangle different kinds of G4 DNA structures *in vitro* with higher catalytic efficiency, as compared to other DNA helicases (Bharti *et al*, 2013). Of note, G4 *foci* were reported to accumulate in FANCJ-deficient human cells, revealing that other G4 resolvases only partially compensate for FANCJ loss (Henderson *et al*, 2014; Summers *et al*, 2021). Recent studies based on multi-color single-molecule localization microscopy have provided direct evidence that in human cells G4 are formed at DNA replication forks behind the CMG complex (either on the leading or lagging strand) and resolved by the combined action of FANCJ and replication protein A (RPA) (Odermatt *et al*, 2020; Lee *et al*, 2021). In recent structural studies of the human CMG complex bound to AND-1 (Rzechorzek *et al*, 2020) or the whole human core replisome (Jones *et al*, 2021), the SepB trimer appears to be located near-perpendicular to the N-tier face of the MCM2-7 complex, where it interfaces with CDC45 and GINS *via* its β-propeller domain, while the SepB α-helix bundle and the C-terminal HMG subdomain project away from the CMG, likely in close proximity to the lagging strand. Thus, FANCJ, bound to the SepB helical bundle, occupies a position in the replisome favorable to promptly hop on the lagging strand between the Okazaki fragments and/or the unwound leading template as it emerges from the MCM complex, where G4 DNA structures are expected to arise more frequently. Notably, in a study carried out in DT40 cells it was suggested that G4 occurring at the replication forks could be resolved by the interplay of DDX11 and its binding partner TIMELESS, a component of the fork-protection complex (Lerner *et al*, 2020). In fact, in this cell system DDX11 and TIMELESS were found to act epistatically in preventing G4-dependent epigenetic instability, but independently from FANCJ. In agreement with these findings, herein we propose the existence of at least two partially redundant pathways responsible for G4 resolution at the DNA replication forks in human cells: one operating on the DDX11-TIMELESS axis and the other one involving FANCJ and its binding partner AND-1 (Calì *et al*, 2016; Cortone *et al*, 2018; Lerner *et al*, 2020). Furthermore, our data suggest that the FANCJ/AND-1 interplay could be also relevant for repairing DNA ICLs. Integrity of the Fanconi anemia pathway is required to fix these DNA lesions and allow restart of stalled DNA replication forks. In a very recent work, it has been demonstrated that ATR-mediated phosphorylation of AND-1 at Threonine 826 (an amino-acid residue of the SepB helical bundle) promotes recruitment of the FANCM/FAAP24 complex at ICL-stalled DNA replication forks (Zhang *et al*, 2022). Of note, the authors of this study have proposed that AND-1 initiates the Fanconi anemia pathway by sensing and specifically binding DNA ICLs through its C-terminal HMG domain: Threonine 826 phosphorylation would elicit an AND-1 conformational change that enhances affinity of the HMG domain for DNA ICLs promoting FANCM/FAAP24 recruitment to these damaged sites.

Our study also addresses the possible functional significance of five *FANCJ* missense variants, that are found in different cancer types and resulting in mutations of CIP box amino-acid residues (Moyer *et al*, 2020; Calvo *et al*, 2021). We found that AND-1-binding by these cancer-related FANCJ mutants is severely compromised, while the DNA helicase activity of at least two of them is not impaired, as compared to the wild type protein. Since our results revealed that association of FANCJ to the DNA replication machinery *via* AND-1 was crucial for preserving genomic integrity, in either in normal or perturbed conditions, the above clinically relevant FANCJ CIP box mutations could be interpreted as cancer risk factors. Further analysis, evaluating tumor frequency in *FANCJ*-knocked-in mouse models is required to assess impact and prognostic relevance of the above cancer-associated *FANCJ* variants.

## Methods

### Plasmid construction and protein expression and purification

Plasmid pCSII-MCS-EF (version 3.4) harboring the human Flag-tagged FANCJ open reading frame (ORF) was a gift from Hisao Masai (Tokyo Metropolitan Institute of Medical Science, Tokyo, Japan) (Uno *et al*, 2012). The FANCJ mutant alleles, named AALA and N734H, D736H, Q740H, Y743C and A745T, were produced by a PCR-mediated site-directed mutagenesis protocol(Carey *et al*, 2013) using the Q5 DNA polymerase (New England Biolabs Laboratories) or KOD Hot start (Merck). Plasmids (pCSII-MCS-EF version 3.4) expressing human Flag-tagged FANCJ wild type (WT) and its mutant forms, named AALA, N734H, D736H, were transfected into HEK 293T cells grown on 10 15-cm dishes. After 48h, cells were detached using ice-cold PBS (150 mM NaCl, 2.7 mM KCl, 10 mM Na_2_HPO4, 1.8 mM KH_2_PO4, pH 7.5) and collected by centrifugation (250 *g* × 3 min). Cell pellets were re-suspended in Binding Buffer (50 mM Tris-HCl, pH 7, 150 mM NaCl, 10% glycerol, 0.1% Triton X-100 and 2 x cOmplete EDTA-free protease inhibitor cocktail [Roche]). Cell lysis was obtained by a mechanical-pestle and sonication, following digestion with Benzonase (Merck) at 25 U/mL (in the presence of 5 mM MgCl_2_) for 30 min. The sample was ultra-centrifuged (21500 *g* × 45 min), the supernatant filtered with 0.22 μM filter and mixed with anti-Flag M2 Agarose beads (Merck) for 2 h in a rotating wheel at 4 °C. Resin was washed with Binding Buffer containing increasing amounts of NaCl (150 mM, 300 mM and 500 mM). Bound protein was eluted from the resin using Elution Buffer (50 mM Tris-HCl, pH 7, 300 mM NaCl, 10% glycerol, 0.1% Triton X-100, 0.2 μg/μL 3x Flag-peptide, 2x cOmplete EDTA-free protease inhibitor cocktail [Roche]). Eluted sample was concentrated on a Vivaspin (cut-off 10 kDa). Sample buffer was exchanged by diafiltration in Storage Buffer (25 mM Tris-HCl, pH: 7.5, 150 mM NaCl, 10% glycerol, 1 mM PMSF and 1 mM dithiothreitol).

Recombinant human 6xHis-tagged AND-1 SepB (amino-acid residues 336-826) was produced in *Escherichia coli* BL21 (*DE3*) cells transformed with pRSF-Duet-1 plasmid construct (a gift from Luca Pellegrini, Cambridge, United Kingdom) and purified, as previously described (Kilkenny *et al*, 2017).

The FANCJ-CIP box (amino-acid residues 730-747) wild type (WT) and AALA mutant were produced as GST-fused proteins by cloning the encoding ORF sequences into the pGEX-6P-1 plasmid (GenScript).

The SepB α-helix-bundle (amino-acid residues 718 – 824) was produced as a GST-fused by cloning the encoding ORF into the pGEX-2T plasmid. The recombinant chimeric protein was expressed in *E. coli* Rosetta p*LysS* (*DE3*) cells, grown in LB medium at 37 °C. When the A_600_ of the culture reached 0.6 optical density (OD), the cell culture was cooled down at 20 °C and isopropyl thio-D-galactopyranoside (IPTG) was added at 0.1 mM. After an overnight incubation at 20 °C, cells were collected by centrifugation and re-suspended in Binding Buffer (PBS containing: 2 x cOmplete EDTA-free protease inhibitor cocktail [Roche], 1 mg/mL Lysozyme, 25 U/mL Turbonuclease and 5 mM MgCl_2_). Cells were broken by sonication and incubated for 20 min on ice, in a shaking platform, to allow further digestion of nucleic acids by Turbonuclease. Thereafter, NaCl concentration was adjusted to 300 mM and the sample was ultracentrifuged (21500 *g* × 45 min). The supernatant was filtered through a 0.22 μM filter and mixed with Glutathione Sepharose 4B agarose beads (Cytiva). The sample was incubated for 1 h in a rotating wheel at 4 °C. Resin was washed with Binding Buffer (PBS [150 mM NaCl, 2.7 mM KCl, 10 mM Na_2_HPO4, 1.8 mM KH_2_PO4, pH 7.5] supplemented with 150 mM NaCl) and protein was eluted with Elution Buffer (50 mM Tris-HCl pH 7.5, 300 mM NaCl and 10 mM reduced glutathione). Eluted sample was concentrated using a Vivaspin system (cut-off 10 kDa). Sample buffer was exchanged by diafiltration in Storage Buffer (50 mM Tris-HCl, pH: 7.5, 300 mM NaCl, 1 mM PMSF and 1 mM dithiothreitol).

All proteins were stored in aliquots at −80 °C.

### Co-immunoprecipitation experiments from whole cell extract

pCSII-EF-MCS- plasmids expressing Flag-tagged FANCJ WT and AALA mutant were transfected into HEK 293T (shown in Figure 1C and 5A) or *FANCJ*-KO HeLa cells (shown in Figure 7B) using poly-ethylenimine (PEI, Polyscience, Inc.). At 48 hr *post*-transfection, cells (about 1 × 10^9^ cells/experiment) were detached, collected by centrifugation (250 *g* × 3 min). After two washes in ice-cold PBS, cell pellets were re-suspended in Lysis Buffer (50 mM Tris-HCl, pH 8.0, 150 mM NaCl, 0.25% [v:v] Triton X-100, 10% [v:v] glycerol) supplemented with cOmplete EDTA-free protease inhibitor cocktail (Roche). The samples were subjected to sonication on ice (8 cycles consisting of 2-s impulses at an output 15% followed by 5-s intervals) and centrifuged for 10 min at 13000 *g* at 4 °C. Then, 20 μL of Flag-M2 agarose beads (Merck) were added to cell extract aliquots containing 1-2 mg of total protein. Samples were incubated at 4 °C for 2 hr in a rotating wheel. Beads were finally washed four times with Lysis Buffer and pull-down proteins were eluted by adding SDS-PAGE loading buffer 2 × (100 mM Tris-HCl, pH 6.8, 20% [v:v] glycerol, 400 mM β-mercapto-ethanol, 1.0% [w:v] SDS, 0.02% [w:v] blue bromophenol) to each pelleted resin sample. Mixtures were incubated at 95 °C for 5 min and subjected to Western blot analysis using the indicated antibodies.

### In vitro pull-down assays

Direct interaction between recombinant purified human AND-1 SepB and FANCJ (shown in Figure 2B and Appendix Figure S1B) proteins was analyzed by co-immunoprecipitation experiments. 10 μL of Flag-M2 agarose beads (Merck) were incubated with Flag-tagged FANCJ wild type or AALA mutant (2 μg of each purified recombinant protein) in mixtures (final volume: 300 μL) containing Pull-down Buffer 1 (20 mM HEPES-NaOH pH 7.2, 150 mM NaCl, 5% [v:v] glycerol, 1 mM dithiothreitol, 0.1% [v:v] Igepal) supplemented with cOmplete EDTA-free protease inhibitor cocktail (Roche). Samples were incubated for 1 hour at 4 °C in a rotating wheel. Then, they were washed 3 times with Pull-down Buffer 1 containing 1% (w:v) Bovine Serum Albumin (BSA) and incubated in the same buffer (final volume: 200 μL) for 20 min at 4 °C in a rotating wheel. Purified AND-1 SepB (4 μg) was added to the indicated samples in Pull-down Buffer 1 containing 1% (w:v) BSA supplemented with cOmplete EDTA-free protease inhibitor cocktail (Roche) (final volume: 250 μL). Samples were incubated for 1 h at 4 °C in a rotating wheel. Then, samples were washed twice with Pull-down Buffer 1 containing 1% (w:v) BSA and protease inhibitor cocktail and four times with Pull-down Buffer 1. Pulled-down proteins were eluted were eluted by adding SDS-PAGE loading buffer 2x (100 mM Tris-HCl, pH 6.8, 20% [v:v] glycerol, 400 mM β-mercapto-ethanol, 1.0% [w:v] SDS, 0.02% [w:v] blue bromophenol) to each pelleted resin sample. Mixtures were incubated at 95 °C for 5 min and subjected to Western blot analysis using the indicated antibodies.

GST-pull-down assay to examine the interaction between GST-FANCJ-CIP box and the AND-1 SepB (shown in Figure 2C) were carried out essentially as previously described(Villa *et al*, 2016). Transformed cells were cultured in 25 mL LB medium at 37 °C until the A_600_ of the culture reached 0.9 - 1.0 optical density (OD). The expression of the fusion protein was induced by adding IPTG at 0.5 mM to the medium. Cell culture was incubated for 20 hrs at 16 °C. Cells were centrifuged at 3500 *g* (30 minutes, at 4 °C), the pellet was resuspended Lysis Buffer (50 mM Tris-HCl, pH 7.0, 500 mM NaCl, 10% [v:v] glycerol, 1 mM dithiothreitol) complemented with cOmplete EDTA-free protease inhibitor cocktail (Roche). After cell disruption by sonication, the cell lysate was clarified by centrifugation at 30000 *g* at 4 °C for 1 hr. The soluble extract was then mixed with Glutathione Sepharose 4B beads (50 μL; Cytiva) pre-equilibrated in the same buffer and incubated under rotation at 4 °C for 1 hr. Unbound protein was removed by three consecutive washes with Lysis Buffer (1 ml), followed by three washes (1 ml) with Pull-down Buffer 1 containing BSA at 1% [w:v]. Then, the purified AND-1 SepB protein (0.5 mg in 500 μL) was added to the beads and the binding was allowed to take place by incubating the mixtures for 1 hr at 4 °C under rotation. Thereafter, the beads were washed three times with Pull-down Buffer 1 and washed again three times with the same buffer without BSA. Proteins were eluted by adding SDS-PAGE loading buffer 2 x to each pelleted resin sample. Mixtures were incubated at 95 °C for 5 min and subjected to SDS-PAGE followed by Coomassie staining.

GST-pull-down assays (reported in Figure 2E and 7C) were carried out to examine the interaction between GST-SepB α-helix bundle and full-length FANCJ WT and AALA. Glutathione Sepharose 4B beads (10 μL) were incubated with GST-SepB α-helix bundle (0.5 μg) in mixtures containing Pull-down Buffer 2 (25 mM Tris-HCl, pH 7.5, 150 mM NaCl and 0.1% [v:v] Triton X-100). Samples were incubated for 1h at 4 °C in a rotating wheel, washed three times with Pull-down Buffer 2. Then, purified Flag-tagged FANCJ WT and mutant forms were added (0.5 μg) to the mixtures containing the beads. Samples were incubated for 1h at 4 °C under rotation and finally washed five times with Pull-down Buffer 2 containing NaCl at 300 mM and Triton X-100 at 1% (v:v). Pulled-down proteins were eluted by adding SDS-PAGE loading buffer 2 x to each resin pellet. Mixtures were incubated at 95 °C for 5 min and subjected to SDS-PAGE followed by Coomassie staining.

### Generation of a FANCJ-knockout (KO) HeLa cell line by CRISPR/Cas9

To establish a FJ-KO line, HeLa cells (ATCCCCL-2) were transiently transfected with SpCas9-expressing PX459 plasmid (Addgene # 62988) using Lipofectamine 2000 (ThermoFisher Scientific). 48 hours *post*-transfection, Alt-R synthetic paired guide RNAs (Integrated DNA Technologies), targeting *FANCJ* exon 2 (TATAAAGCTTACCCGTCACA) and exon 3 (TGTTTGTTGGAGAGTCCCAC), were transfected using Lipofectamine RNAiMAX Transfection Reagent (ThermoFisher Scientific) and expanded for clonal populations. These latter were screened using Western blot analysis to detect FANCJ protein expression and further confirmed by Sanger sequencing. For sequencing, *FANCJ* genomic *loci* were amplified using high-fidelity Q5 DNA polymerase (New England Biolabs) and amplified fragments were subcloned using the Zero Blunt Topo PCR cloning kit (Invitrogen). Plasmid DNA was isolated from at least 10-15 colonies and sequenced to identify frameshifts or deletions produced by the Cas9 activity. *CRISPRscan* (https://www.crisprscan.org/) and *CRISPOR* (http://crispor.tefor.net/) online tools were used to design guide RNAs.

### Cell viability assays

For cell viability assays, HeLa wild type control (Ctrl) and *FANCJ*-KO cells were seeded at a concentration of 1000-1500 cells/well in 96-well plates. Briefly, cells were treated with the indicated concentrations of PDS and MMC for 5-6 days (chronic treatment). Cell viability was analyzed using a crystal violet solution (0.5% [w:v] dissolved in 20% (v:v) methanol).

### Establishment of stable FJ-KO HeLa cell lines expressing FANCJ WT and the AALA mutant

For establishing FJ-KO HeLa cell lines that express FANCJ WT or the AALA variant, HeLa FJ-KO cells were transfected with the following plasmids: pMK204-TetOne AAVS1-MCS (+) Flag-tagged FANCJ WT or pMK240-TetOne AAVS1-MCS (+) Flag-tagged FANCJ-AALA (GenScript). The plasmid pMK240-TetOne AAVS1-MCS (+), was a gift from Masato Kanemaki (National Institute of Genetics, Mishima, Japan) (Okumura *et al*, 2018). After transfection, cells were cultured in 96-well plates and selected by adding Puromycin (0.3 μg/mL) to DMEM complete medium. Single clones were isolated and further expanded in the presence of Puromycin at a lower concentration (0.2 μg/mL). Expression of Flag-tagged FANCJ, WT and AALA mutant, was assessed before and after induction with Doxycycline (1 μg/mL) by Western blot analysis using an anti-FANCJ (cat. B1310, Merck) and/or an anti-Flag antibody.

All cell lines were tested for mycoplasma contamination and maintained at 37 °C with 5% CO_2_.

### Co-immunoprecipitation experiments from cell nuclear fraction

Co-immunoprecipitations were carried out on nuclear extracts prepared from HEK 293T cells (about 4 × 10^7^ cells/experiment), transiently transfected with the indicated plasmid constructs. After transfection, cell cultures were synchronized in S phase with a single block in thymidine (2 mM) followed by release in fresh medium for 2.5 hr. Preparation of cell nuclear fraction was according to a published protocol with modifications(Guillou *et al*, 2010). Cell pellets were re-suspended in 1 mL of Osmotic Buffer (10 mM Hepes-NaOH, pH 7.9, 0.2 M potassium acetate, 0.34 M sucrose, 10% [v:v] glycerol, 1 mM dithiothreitol, 0.1% [v:v] Triton X-100) and incubated for 5 min on ice. After centrifugation (800 *g* for 5 min), the nucleus/chromatin fraction present in the pellet was re-suspended in 1 mL of Hypotonic Buffer (10 mM Hepes-NaOH, pH 7.9, 50 mM NaCl, 1 mM dithiothreitol, 0.1% [v:v] Triton X-100) supplemented with cOmplete EDTA-free protease inhibitor cocktail (Roche). Samples were subjected to sonication on ice (10 cycles consisting of 10-s impulses at an output 10% followed by 20-s intervals) followed by incubation for 20 min at 37 °C in the presence of micrococcal nuclease (2 units/sample; Merck, cat. # N3755) and CaCl_2_ (at 10 mM). Insoluble material was removed by centrifugation at 16,000 *g* for 30 min. Samples (containing 0.3-0.5 mg of protein) were used in immunoprecipitation experiments with the anti-Cdc45 rabbit antibodies and control rabbit IgG bound to Protein A Sepharose beads (GE Healthcare). Mixtures were incubated for a minimum of 3 hr (or overnight) at 4 °C in a rotating wheel. Beads were washed 4 times with Washing Buffer (10 mM Hepes-NaOH, pH 8.0, 50 mM NaCl, 1 mM dithiothreitol, 0.1% [v:v] Triton X-100). Proteins bound to the beads were eluted with SDS-PAGE loading buffer, incubated at 95 °C for 5 min and analyzed by Western blot using the indicated antibodies.

### SiRF assay

Quantitative *in situ* analysis of protein interactions at DNA replication forks (SiRF) assay was carried out as previously described (Roy *et al*, 2018). HeLa cells (wild type control, FJ-KO and stably complemented cell lines) were incubated with Doxycycline (1 μg/mL) for 24h before seeding on 8-well chamber slides (Lab-Tek), at approximately 15000 cells/well. Cells were pulsed with EdU (125 μM) for 15 min, followed by an incubation with hydroxyurea (HU, 4 mM) for stalling DNA replication forks. Cells were subjected to fixation with a solution containing 2% (v:v) paraformaldehyde in PBS for 15 min at room temperature and permeabilization with a solution containing 0.25% (v:v) Triton X-100 in PBS for 15 min at RT, followed by washing with PBS. EdU-biotinylation was performed with a click-it reaction cocktail (2 mM CuSO_4_, 10 μM biotin-azide, 100 mM sodium ascorbate in PBS). Slides were incubated with primary antibodies, mouse anti-biotin (clone BTN.4, ThermoFisher Scientific) and rabbit anti-FANCJ (cat. B1310, Merck). Specific probes that cross-react with the primary antibodies were used for rolling circle amplification and the related SiRF signals were visualized as discrete PLA (proximity ligation assay) spots in cell nuclei that were also counter-stained with DAPI. Images were acquired with a confocal microscope (Zeiss LSM 700) using a 63x magnification objective (Nikon). Signals were quantified using ImageJ software (1.52v) and the Find Maxima tool with variable values of prominence depending on each experiment. Each cell was analyzed and quantification (number of points identified), done using the software, was also verified by eye inspection. Control experiments were also carried out to detect sites of DNA synthesis in cell nuclei by EdU-EdU PLA spot analysis for the same aforementioned lines.

### Immunofluorescence

For analyzing γ-H2A.X *focus f*ormation, HeLa cells (wild type control, FJ-KO and complemented cell lines) were grown overnight on coverslips and induced with Doxycycline (1 μg/mL) for 24h and further treated for 24h with PDS (1 μM) or MMC (10 nM). After incubation, coverslips were fixed in cold methanol for 5 min and further blocked/permeabilized for 1h at room temperature with PBS containing: 1% (w:v) bovine serum albumin (BSA), 0.3 M glycine, 0.1% [v:v] Tween-20. Coverslips were incubated overnight with rabbit monoclonal anti-γ-H2A.X (phospho S139; 1:250, clone EP854(2)Y, Abcam) in a humid chamber at 4 °C. After washing with PBS, coverslips were incubated for 1h at room temperature with goat-anti-rabbit Alexa Fluor™ 488 (1:500, ThermoFisher Scientific) as a secondary antibody. Thereafter, coverslips were mounted into glass slides using mounting media containing DAPI. Images were acquired with a confocal microscope (Zeiss LSM 700) using a 63x magnification objective (Nikon). γ-H2A.X *foci* were quantified with ImageJ (1.52v) using the Find Maxima tool with variable values of prominence depending on each experiment: each cell was analyzed, and quantification (number of *foci* identified) made by the software was also checked and confirmed by eye inspection.

### DNA fiber track assays

Cells were pulse-labeled with chloro-deoxyuridine (CldU, 20 μM) for 30 min, followed by incubation in iodo-deoxyuridine (IdU, 200 μM) for 30 min. For the experiments where a PDS treatment was used, the drug was administered at 1 μM to the cells together with IdU. Approximately 1750 cells were lysed with 7.5 μL of Lysis Buffer (200 mM Tris-HCl, pH 7.5, 50 mM EDTA, 0.5% [v:v] SDS). Fibers were spread on Superfrost microscope slides (Epredia, AG00008332E01MNZ10), which were tilted by ~45° and air-dried for 2 min. Slides were then fixed in a mixture of methanol:acetic acid (3:1, v:v). After drying, or on the next day, DNA was denatured using a solution of HCl (2.5 M) for 60 min. After a careful wash with PBS, slides were incubated with filtered Blocking Solution (PBS containing 1% [w:v] BSA) for 60 min at room temperature. Slides were incubated with a rat anti-BrdU antibody (1:200, clone BU1/75 (ICR1), Abcam) overnight at 4 °C. Next day, slides were washed with PBS and incubated with goat anti-rat antibody labelled with Alexa Fluor™ 647 (1:500, ThermoFisher Scientific). Thereafter, slides were incubated with a mouse anti-BrdU antibody (1:500, clone B44, BD Biosciences), followed by incubation with a goat anti-mouse antibody labelled with Alexa Fluor™ 555 (1:500, ThermoFisher Scientific), both at room temperature for 1 h. Slides were mounted with mounting media containing DAPI (PBS:glycerol, 1:1 [v:v]). Images of DNA fibers were taken with a confocal microscope (Zeiss LSM 700) using a 63x magnification objective (Nikon). Fiber tract lengths were assessed with ImageJ and μm values were converted into DNA kilobases using the conversion factor 1 μm = 2.59 kb for the spreading technique (Quinet *et al*, 2017; Nieminuszczy *et al*, 2016). Sister forks derived from the same fiber were also measured for fork asymmetry analysis.

### DNA helicase assay

PAGE-purified oligonucleotides used for the preparation of DNA substrates were purchased from Merck. An anti-parallel bi-molecular G4 DNA substrate was prepared using the following fluorescent-labelled DNA oligonucleotide OX1 (5’-GACCACTG-[Cy3]T-CGGTTCCAAGCACTGTCGTACTTGATATTTTGGGGTTTTGGGG-3’), as previously described(Calì *et al*, 2016). DNA helicase assays were carried out in reaction mixtures (20 μL) containing the indicated proteins in buffer 25 mM Hepes-NaOH, pH 7.5, 5 mM MgCl_2_, 25 mM KCl, 2 mM dithiothreitol, 0.1 mg/mL [w:v] BSA, 2 mM ATP, 7.5 nM DNA substrate. Reactions were initiated by addition of the indicated proteins and then incubated for 20 min at 37 °C. Reactions were quenched with the addition of 5 μl of 5 x Stop Solution (0.5% [w:v] SDS, 40 mM EDTA, 0.5 [w:v] mg/mL proteinase K, 20% [v:v] glycerol). Samples were run on a 8% polyacrylamide-bis (29:1) gel in TBE containing 0.1% [w:v] SDS at a constant voltage of 100 V. Both gel and running buffer contained KCl at 10 mM to preserve G4 DNA structure. After electrophoresis, gels were analyzed using an imaging system instrument (ChemiDoc, Bio-Rad Laboratories). Displaced oligonucleotide was quantified and any free oligonucleotide in absence of enzyme was subtracted.

### Statistical analysis

Statistical significance (*p* values), analysis and graphs were performed using GraphPad prism 6 software. Statistical tests and differences are mentioned in the respective figure legends.

## Acknowledgements

The authors acknowledge Hisao Masai (Tokyo Metropolitan Institute of Medical Science, Tokyo, Japan) for the pCSII-EF-MCS plasmid expressing Flag-tagged FANCJ WT; Masato Kanemaki (National Institute of Genetics, Mishima, Japan) for the plasmid pMK240-TetOne AAVS1-MCS (+); Luca Pellegrini (University of Cambridge, Cambridge, United Kingdom) for the plasmid pRSF-Duet-1 expressing 6xHis SepB. This study has received funding from the European Union’s Horizon 2020 research and innovation programme under the Marie Skłodowska-Curie grant agreement n. 859853 (acronym: AntiHelix; to A.B., D.S., F.M.P., L.M.R.N. and S.O.), and grant agreement n. 722729 (acronym: Syntrain; to N.K.J. and D.B.), from the Cross-border Cooperation Programme Italy-Slovenia 2014–2021 by the European Regional Development Fund and national funds (TransGlioma), and from *Associazione Italiana per la Ricerca sul Cancro* (AIRC IG n. 25965 to F.M.P.; AIRC IG 23710 and IG 18976 to D.B.; AIRC IG 20778 to S. O.).

## Author contributions

A.B. designed and carried out most of the experiments; D.S. purified the AND-1 6xHis-SepB; S.O. produced the FANCJ and FANCJ CIP/AND-1 homology model; L.M.R.N. performed the GST-pull-down assays shown in Figure 2D and the co-immunoprecipitation assays reported in Appendix Figure S1A; G.C. produced the pCSII-EF-MCS plasmid expressing Flag-tagged FANCJ AALA mutant and carried out some of the co-immunoprecipitations in cell extracts; N.K.J. generated the FJ-KO HeLa cell line and did the cell viability assays; A.B., L.M.R.N., S.O., N.K.J., D.B. and F.M.P. analyzed the data; F.M.P. identified the FANCJ CIP box, conceptualized and supervised the project and wrote the manuscript with the help of A.B. All the authors read and approved the manuscript.

## Materials and Correspondence

Material requests and correspondence should be addressed to F.M.P.

